# *Topaz1*, an essential gene for murine spermatogenesis, down-regulates the expression of numerous testis-specific long non-coding RNAs

**DOI:** 10.1101/2021.03.09.434544

**Authors:** Manon Chadourne, Elodie Poumerol, Luc Jouneau, Bruno Passet, Johan Castille, Eli Sellem, Eric Pailhoux, Béatrice Mandon-Pépin

**Affiliations:** Université Paris-Saclay, UVSQ, INRAE, BREED, 78350, Jouy-en-Josas, France; Ecole Nationale Vétérinaire d’Alfort, BREED, 94700, Maisons-Alfort, France; Université Paris Saclay, INRAE, AgroParisTech, GABI, Jouy-en-Josas, France; R&D Department, ALLICE, Paris, France

## Abstract

Spermatogenesis involves coordinated processes, including meiosis, to produce functional gametes. We previously reported *Topaz1* as a germ cell-specific gene highly conserved in vertebrates. *Topaz1* knockout males are sterile with testes that lack haploid germ cells because of meiotic arrest after prophase I. To better characterize *Topaz1^−/−^* testes, we used RNA-sequencing analyses at two different developmental stages (P16 and P18). The absence of TOPAZ1 disturbed the expression of genes involved in microtubule and/or cilium mobility, which was consistent with testicular histology showing the disruption of microtubules and centrosomes. Moreover, a quarter of P18 dysregulated genes are long non-coding RNAs (lncRNAs), and three of them are testis-specific and located in spermatocytes, their expression starting between P11 and P15. The suppression of one of them, *4939463O16Rik*, did not alter fertility although sperm parameters were disturbed and sperm concentration fell. The transcriptome of P18-*4939463O16Rik*^−/−^ testes was altered and the molecular pathways affected included microtubule-based processes, the regulation of cilium movement and spermatogenesis. The absence of TOPAZ1 protein or *4930463O16Rik* produced the same enrichment clusters in mutant testes despite a contrasted phenotype on male fertility. In conclusion, TOPAZ1 appeared to stabilize the expression of numerous lncRNAs. Its suppression is not essential for fertility but required during the terminal differentiation of male gametes.

**Author Summary:** The *Topaz1* gene was initially characterized during the initiation of meiosis in the sheep fetal ovary. In order to determine its function, a KO of the murine gene was performed. In this species, only males were sterile and spermatogenesis was blocked before the first meiotic division. Here, we show that cytoskeletal elements are markedly disturbed in mutant testes, indicating that these elements play an important function in spermatogenesis. While the mitotic spindle of spermatogonia was normal, the meiotic spindle of spermatocytes was hemi-spindle-shaped and the homologous chromosome pairs could position themselves on the equatorial plate. In addition, lncRNAs account for 25% of genes whose expression in testes varies significantly in the absence of *Topaz1*. This suggests a key role for these factors in spermatogenesis. Largely testis-specific, they may be involved in spermatogenesis and play a more or less critical role in mouse fertility, which probably also depends on their redundancies.

## Introduction

In mammals, an organism derives from two parental haploid gametes, a maternal oocyte and paternal sperm. Meiosis is a highly specialized event that leads to the production of these haploid germ cells (Kleckner, 1996). In females, meiosis is initiated during fetal life while male germ cells are involved in the meiosis process around puberty. In males, meiosis is essential during spermatogenesis that involves mitotic division and the multiplication of spermatogonia, the segregation of homologous chromosomes and the spermiogenesis of haploid germ cells. This complex process of spermatogenesis, which progresses through precisely timed and highly organized cycles, is primordial for male fertility. All these different events are highly regulated and associated with the controlled expression of several testis-enriched genes. A previous study had demonstrated the essential role of *Topaz1* during meiosis in male mice (Luangpraseuth-Prosper et al., 2015). *Topaz1* is a highly conserved gene in vertebrates (Baillet et al., 2011). Its expression is germ cell-specific in mice (Baillet et al., 2011). The suppression of *Topaz1* in mice (*Topaz1^−/−^*) results in azoospermia (Luangpraseuth-Prosper et al., 2015). Male meiotic blockage occurs without a deregulation of chromosome alignment and TOPAZ1 is not involved in formation of the XY body or the maintenance of MSCI (Meiotic Sex Chromosome Inactivation). *Topaz1* depletion increases the apoptosis of mouse adult male pachytene cells and triggers chromosome misalignment at the metaphase I plate in mouse testes (Luangpraseuth-Prosper et al., 2015). This misalignment leads to an arrest at the prophase to metaphase transition during the first meiosis division (Luangpraseuth-Prosper et al., 2015). Microarray-based gene expression profiling of *Topaz1^−/−^* mouse testes revealed that TOPAZ1 influences the expression of one hundred transcripts, including several long non-coding RNA (lncRNAs) and unknown genes, at postnatal day 20 (P20) (Luangpraseuth-Prosper et al., 2015).

Since discovery of the maternal *H19* lncRNA (Brannan et al., 1990) and the *Xist* (Brockdorff et al., 1992) genes that regulate the structure of chromosomes and mediate gene repression during X chromosome inactivation, interest in studying the role of non-coding RNAs (ncRNAs) has grown considerably. Non-coding RNAs are present in many organisms, from bacteria to humans, where only 1.2% of the human genome codes for functional proteins (Carninci et al., 2005; ENCODE Project Consortium et al., 2007; Gil and Latorre, 2012). While much remains to be discovered about the functions of ncRNAs and their molecular interactions, accumulated evidence suggests that ncRNAs participate in various biological processes such as cell differentiation, development, proliferation, apoptosis and cancers.

They are divided into two groups: small and long non-coding RNAs (sncRNAs and lncRNAs, respectively). SncRNAs contain transcripts smaller than 200 nucleotides (nt). They include microRNAs (miRNAs, 20-25 nt), small interfering RNAs (siRNAs), PIWI-interacting RNAs (piRNAs, 26-31 nt) and circular RNAs (cricRNA). These sncRNAs are essential for several functions such as the regulation of gene expression and genome protection (Ref in (Morris and Mattick, 2014)) as well as during mammalian spermatogenesis (Yadav and Kotaja, 2014; Bie et al., 2018; Quan and Li, 2018). The second group, lncRNAs, contains transcripts longer than 200 nt without a significant open reading frame. Advances in high-throughput sequencing have enabled the identification of new transcripts, including lncRNAs, most of which are transcribed by RNA polymerase II and possess a 5’ cap and polyadenylated tail (ref in (Jarroux et al., 2017)). They are classified according to their length, location in the genome (e.g. surrounding regulatory elements) or functions.

Several studies have pointed out that testes contain a very high proportion of lncRNA compared to other organs (Necsulea and Kaessmann, 2014; Sarropoulos et al., 2019). However, this high testicular expression is only observed in the adult organ, as the level of lncRNAs in the developing testis is comparable to that seen in somatic organs (Sarropoulos et al., 2019). In mice, some testis-expressed lncRNAs were functionally characterized during spermatogenesis. Thus, the lncRNA *Mrhl* repressed *Wnt* signaling in the Gc1-Spg spermatogonial cell line, suggesting a role in spermatocyte differentiation (Arun et al., 2012). Expression of the Testis-specific X-linked gene was specific to, and highly abundant in, mouse pachytene-stage spermatocytes and could regulate germ cell progression in meiosis (Anguera et al., 2011). Moreover, in male germ cells, it has been shown that the *Dmrt1*-related gene negatively regulates *Dmrt1* (doublesex and mab-3 related transcription factor 1) and that this regulation might be involved in the switch between mitosis and meiosis in spermatogenesis (Zhang et al., 2010). Lastly, in a non-mammalian model as *Drosophila*, Wen *et al*. produced mutant fly lines by deleting 105 testis-specific lncRNAs and demonstrated the essential role of 33 of them in spermatogenesis and/or male fertility (Wen et al., 2016).

Following a previous study, which presented comparative microarray analyses of wild-type and *Topaz1^−/−^* testis RNAs at P15 and P20 (Luangpraseuth-Prosper et al., 2015), we have now performed deep sequencing by bulk RNA-sequencing (RNA-seq) of these testes collected at P16 and P18 in an aim to refine the developmental stages that display transcriptional differences between the two mouse lines. Since the proportion of deregulated lncRNAs represented about a quarter of the differentially expressed genes (DEGs), we studied the testicular localization of three of them. In order to approach the role of testicular lncRNAs, we created a mouse line in which one of them was deleted (*4930463O16Rik*). These knockout mice displayed normal fertility in both sexes, but the male mutants produced half as much sperm as wild-type controls.

## Results

### *Topaz1* mutant testes have a deregulated transcriptome as early as P16

To expand on the previous comparative microarray analyses of wild-type and mutant testes RNA performed at P15 and P20 during the first wave of spermatogenesis (Luangpraseuth-Prosper et al., 2015), transcriptomic analyses by RNA-seq were performed on WT and *Topaz1^−/−^* mouse testes at two developmental stages: P16 and P18. The P16 stage was chosen because these previous microarray analyses had revealed that only the *Topaz1* gene was expressed differently at P15, its expression starting from 5 d*pp*. This means that TOPAZ1 should have had an effect just after P15. Furthermore, whereas at P15, seminiferous tubules contain spermatocytes that have advanced to mid and late-pachytene, at P16 they contain spermatocyte cells that have progressed from the end-pachytene to early diplotene of meiosis I. At P20, the first round spermatids appear, while late-pachytene spermatocytes are abundant at P18 and the very first spermatocytes II appear (Drumond et al., 2011). Therefore, the P16 and P18 stages chosen for this study surrounded as closely as possible the time lapse just before and after the first meiosis I division of spermatogenesis.

Differential analyses of RNA-seq results revealed that 205 and 2748 genes were significantly deregulated in *Topaz1^−/−^* testes compared to WT at P16 and P18, respectively (adjusted p-value (Benjamini-Hochberg) <0.05 and absolute Log2 Fold Change >1 (Log2FC>1) (Figure 1A, Supplementary Table 1). At P16, out of the 205 DEGs, 97 genes were significantly down-regulated (Log2FC<-1 or FC<0.5) and 108 were up-regulated (Log2FC>1 or FC>2). However, at P18, down-regulated DEGs accounted for 91% (2491 genes) and up-regulated genes for only 9% (257 genes). Among all these DEGs, 120 were common to both P16 and P18 (Figure 1A). According to the mouse gene atlas, the 2748 DEGs at developmental stage P18 were largely testis-enriched DEGs in mouse testis-specific genes (Figure 1B).

**Figure 1:**
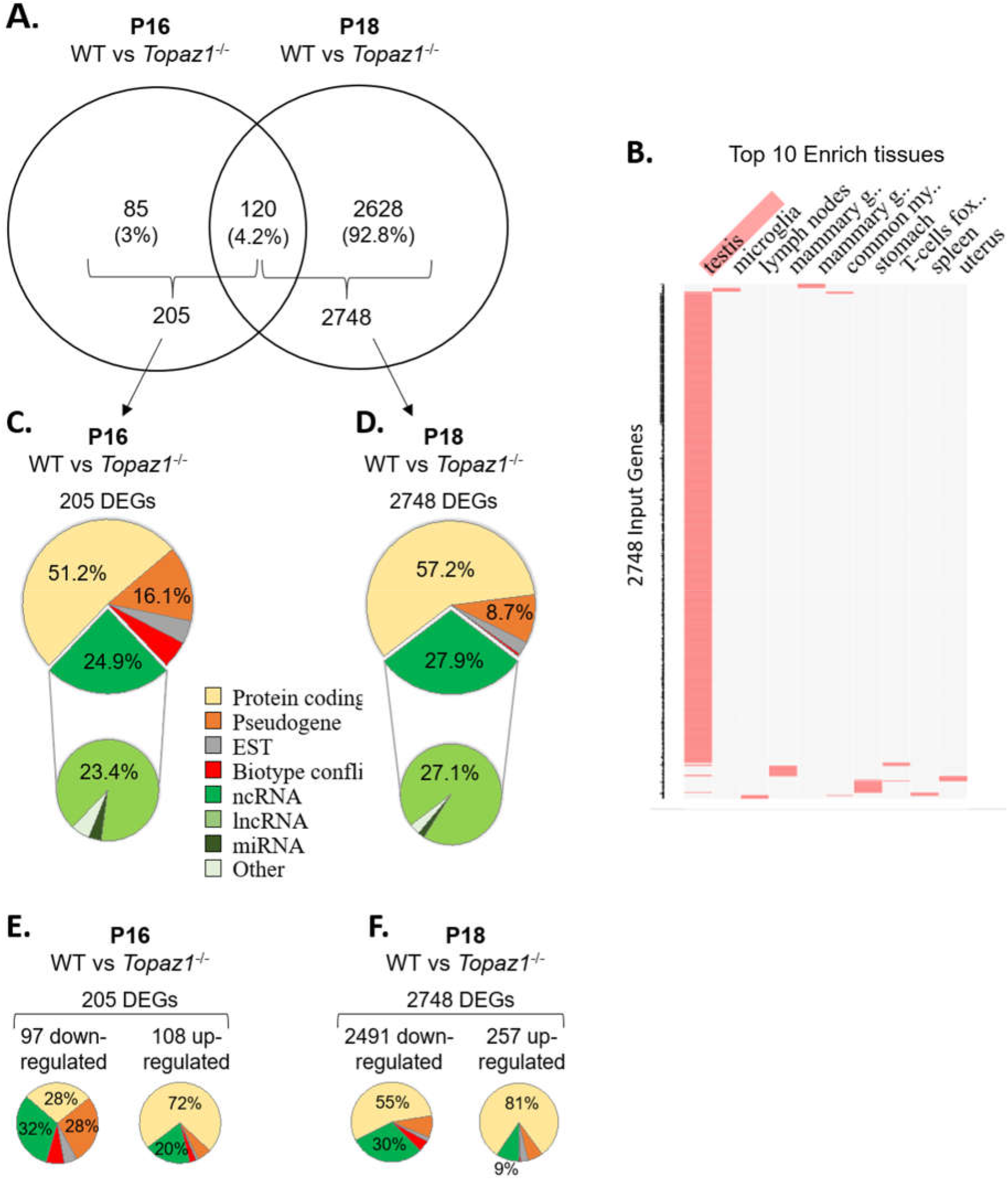
WT vs *Topaz1*^−/−^ deregulated gene analysis from mouse testes. (A) Venn diagram showing the overlap of differentially expressed genes between P16 and P18 *Topaz1^−/−^* mouse testes (adjusted p<0.05 and down-regulated FC<0.5 (log2FC<-1) or up-regulated FC>2 (log2FC>1)). (B) Clustergrammer was generated by the Enrichr website. Top 10 enriched tissues are the columns, input genes (2748 DEGs of P18 *Topaz1^−/−^* compared to normal testes) are the rows, and cells in the matrix indicate whether a DEG is associated with a tissue from the Mouse Gene Atlas. (C-D) Biotype of DEGs in *Topaz1^−/−^* testes at (C) P16 and (D) P18. Around half of them are protein-coding genes whereas around one quarter is ncRNA at both developmental stages. (E-F) Biotype of DEGs in *Topaz1^−/−^* testes at (E) P16 and (F) P18, depending on whether they were up- or down-regulated.

The validation of several DEGs was achieved using RT-qPCR. Two randomly selected genes up-regulated at P16 (B3galt2 and Hp) and three at P18 (*B3galt2*, *Afm* and *Cx3cr1*), four gene down-regulated at P16 and P18 (*Gstt2*, *4930463O16Rik*, *4921513H07Rik* and *Gm21269*) and two non-differential genes (*Cdc25c* and *Nop10*) were analyzed (Supplementary Figure 1). The results confirmed those obtained using RNA-seq.

The biotypes of the differential transcripts (protein-coding, non-coding RNAs, etc.) were determined from the annotations of the NCBI, MGI and Ensembl databases. Two major deregulated groups were highlighted at both stages. The protein-coding gene biotype accounted for half of the deregulated genes (51.2% and 57.2% at P16 and P18, respectively) (Figure 1C-D). A quarter of *Topaz1^−/−^* DEGs; 24.9% and 27.9% at P16 and P18, respectively, was found to belong to the second ncRNA group. Among the latter, the major biotype was lncRNAs at both stages, being 23.4% and 27.1% at P16 and P18, respectively. This significant proportion of deregulated lncRNA thus raised the question of their potential involvement in spermatogenesis.

### Pathway and functional analysis of DEGs

To further understand the biological functions and pathways involved, these DEGs were functionally annotated based on GO terms and KEGG pathway or on InterPro databases through the ontological Database for Annotation, Visualization and Integrated Discovery (DAVID v6.8, https://david.ncifcrf.gov/) using the default criteria (Huang et al., 2009a, 2009b).

At P16, so therefore before the first meiosis division, out of 205 differentially expressed genes, 32% of down-regulated and 20% of up-regulated genes corresponded to non-coding RNAs with no GO annotation or no pathway affiliation for the vast majority (Figure 1E), leading to less powerful functional annotation clustering (Supplementary Table 2). Five clusters with an enrichment score >1.3 were obtained (an enrichment score >1.3 was used for a cluster to be statistically significant, as recommended by Huang et al., (Huang et al., 2009a) but the number of genes in each cluster was small except for annotation cluster number 4. In this, an absence of TOPAZ1 appeared to affect the extracellular compartment. The others referred to the antioxidant molecular function and the biological detoxification process, suggesting stressful conditions.

At P18, corresponding to the first transitions from prophase to metaphase, and considering either all DEGs (2748 DEGs; 2404 DAVID IDs) or only down-regulated genes (2491 DEGs; 2164 DAVID IDs) in the P18 *Topaz1^−/−^* versus WT testes, it was possible to identify five identical clusters with an enrichment score higher than 12 (Figure 1F, Supplementary Table 3). However, the enrichment scores were higher when only down-regulated genes were considered. These clusters include the following GO terms: (*i*) for cellular components: motile cilium, ciliary part, sperm flagellum, axoneme, acrosomal vesicule; (*ii*) for biological processes: microtubule-based process, spermatogenesis, germ cell development, spermatid differentiation (Supplementary Table 3).

Finally, using the InterPro database, four clusters with enrichment scores >1.3 were obtained based on down-regulated genes (Supplementary Table 3) and with up-regulated genes, an absence of TOPAZ1 from mouse testes highlighted the biological pathway of the response to external stimulus or the defense response in the testes. Once again, and as for P16, this suggested stressful conditions in these *Topaz1^−/−^* testes.

These results indicate that an absence of TOPAZ1 induced alterations to the murine transcriptome of the mutant testis transcriptome as early as 16 days after birth. Two days later (P18), these effects were amplified and predominantly involved a down-regulation of genes (91% of DEGs). The loss of TOPAZ1 appeared to disrupt the regulation of genes involved in microtubule and/or cilium mobility, spermatogenesis and first meiotic division during the prophase to metaphase transition. This was in agreement with the *Topaz1^−/−^* phenotype in testes.

### Absence of TOPAZ1 leads to drastic cytoplasmic defects before the first meiotic division

According to the preceding GO pathway analyses showing that a majority of deregulated proteins are involved in microtubule cytoskeleton organization, microtubule-based movements and processes, microtubule organizing centers, centrosomes and centrioles in P18 *Topaz1^−/−^* testes (Supplementary Table 3), we decided to better characterize the cytoplasmic components of germ cells in *Topaz1^−/−^* testes before the first meiotic division. As the meiotic spindle is a key component of these cells before and during the metaphase stage, we studied it using α- and γ-tubulin immunofluorescence (IF) staining for markers of microtubule spindle and centrosome, respectively (Figure 2). We observed one monopolar spindle centered in the germ cells of the *Topaz1^−/−^* testes. Moreover, centrosome staining was diffuse and weak in these cells. This was also observed on entire seminiferous sections (Supplementary Figure 2). The chromosomes were not aligned along a metaphase plate but adopted an atypical rosette shape (Figure 2), reflecting a marked perturbation of the microtubule and centrosome pathways in *Topaz1*-deficient spermatocytes that could lead to meiotic arrest.

**Figure 2:**
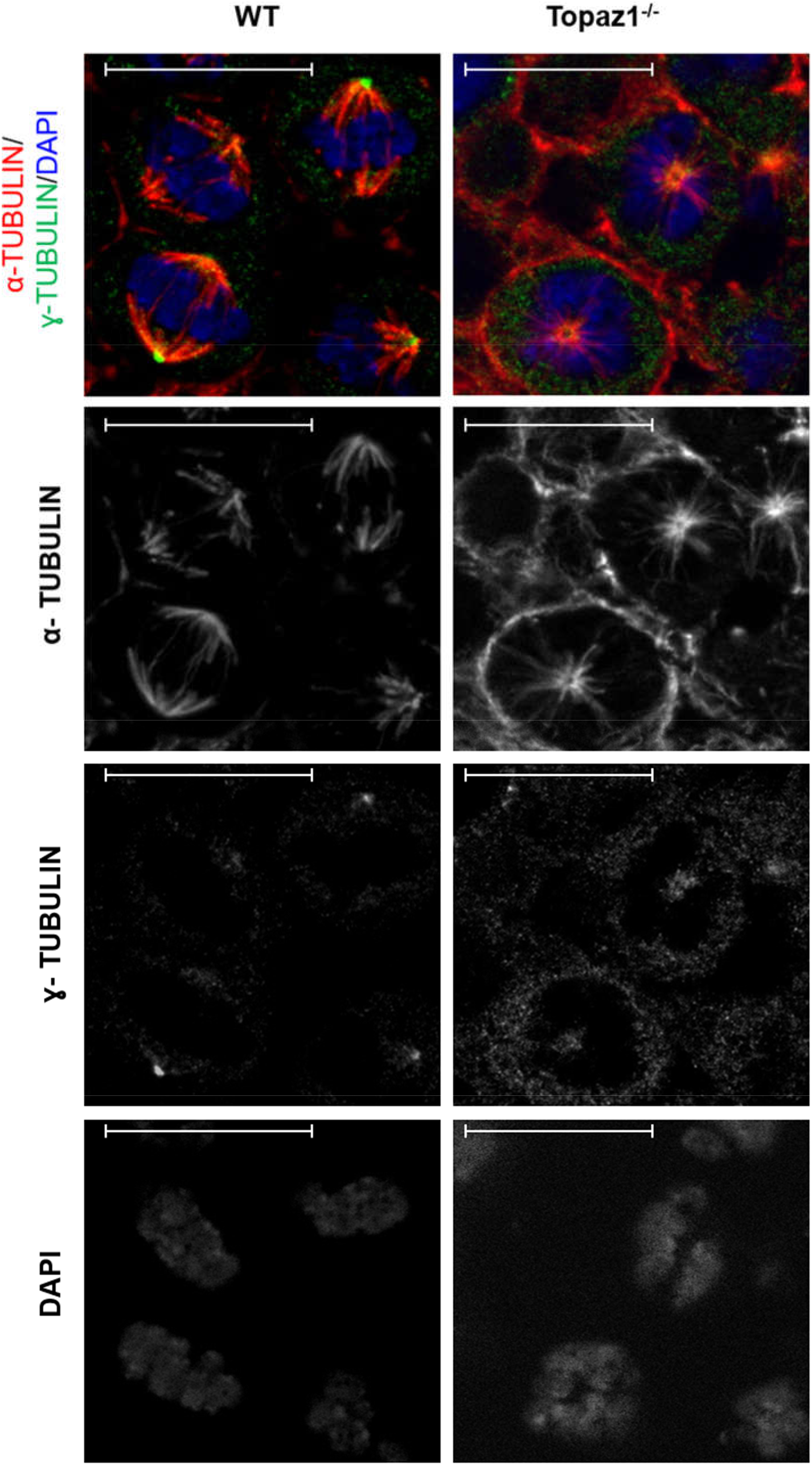
Abnormal metaphase phenotype in *Topaz1*-deficient gonads. Immunofluorescence staining for α-TUBULIN (red), γ-TUBULIN (green) and DAPI (blue) in WT (left) and *Topaz1^−/−^* (right) 28 d*pp* testes sections. Unlike the meiotic metaphases seen in normal testes (left), the metaphases are abnormal in *Topaz1^−/−^* mutants (right) with and atypical rosette shape and hemispindle. Scale bar = 20μm

### Selection of 3 deregulated lncRNA: all spermatocyte-specific

The vast majority of deregulated lncRNAs in *Topaz1^−/−^* testes has an unknown function. We decided to study three of the 35 down-regulated lncRNAs that are shared at the P16 and P18 stages, namely *4930463O16Rik* (ENSMUSG00000020033), *4921513H07Rik* (ENSMUSG00000107042) that is the most down-regulated gene at P16 with a Log2FC of 11.85, and both already highlighted by the previous microarray comparative analyses (Luangpraseuth-Prosper et al., 2015), and *Gm21269* (ENSMUSG00000108448), which has the lowest adjusted p-value at P18. We quantified these transcripts by qPCR in several somatic tissues (brain, heart, liver, lung, small intestine, muscle, spleen, kidney, epididymis and placenta) and in the gonads (testes and ovary). These three lncRNAs were almost exclusively expressed in testes (Figure 3A, C, E). These results were in agreement with RNA-seq data available for *4930463O16Rik* and *Gm21269* on the ReproGenomics viewer (https://rgv.genouest.org/) (Supplementary Figures 3A and 4A, respectively) (Darde et al., 2015, 2019). Our RNA-seq results, summarized using our read density data (bigwig) and the Integrative Genomics Viewer (IGV; http://software.broadinstitute.org/software/igv/), revealed little or no expression of these three genes in *Topaz1^−/−^* testes (Supplementary Figure 5A-C).

**Figure 3:**
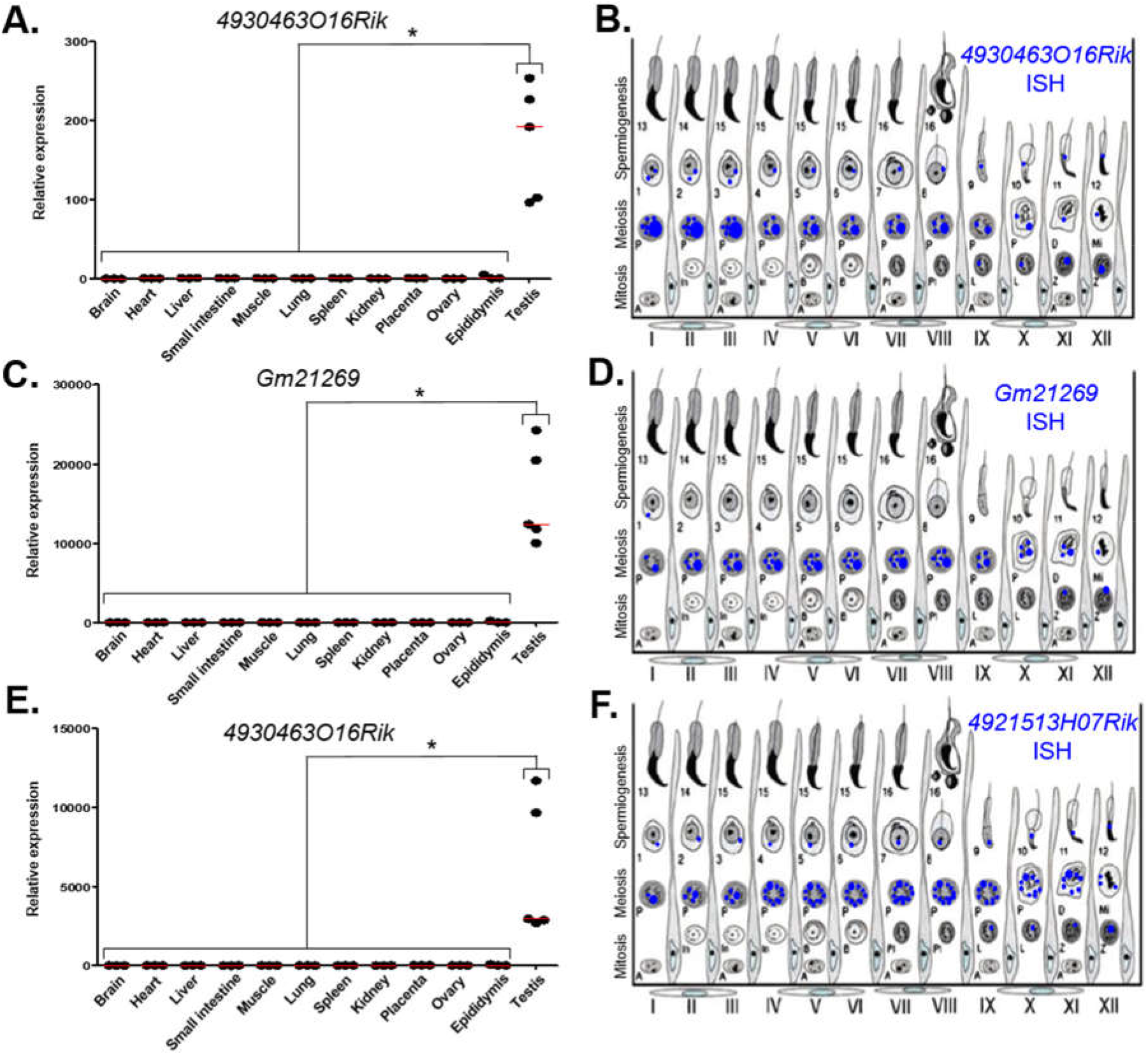
Expression analysis of three lncRNAs. (A-C-E) RT-qPCR analysis of three different lncRNAs. (A) *4930463O16Rik*; (C) *Gm21269*; (E) *4921513H07Rik* in different two month-old tissues from WT mice. The red lines represent the median for each tissue; n=5 for testes and n=3 for other organs. Statistical analyses were performed using the non-parametric Kruskal-Wallis test. * = p<0.05 (B-D-F) Schematic representation of the results of (B) *4930463O16Rik*, (D) *Gm21269* and (F) *49215113H07Rik IS*H expression in meiotic and post-meiotic cells of the WT mouse seminiferous epithelial cycle.

Quantification of these transcripts using qPCR from postnatal to adulthood in WT and *Topaz1^−/−^* testes had previously been reported, as for *4930463O16Rik* and *4921513H07Rik* (Figure 9 in (Luangpraseuth-Prosper et al., 2015)) or performed for *Gm21269* (Supplementary Figure 6, also including the postnatal expression of *4930463O16Rik* and *4921513H07Rik*). The difference in expression between normal and *Topaz1^−/−^* testes was detected as being significant as early as P15 (detected as insignificant in the previous microarray analysis and *Gm21269* was absent from the microarray employed). All showed an absence of expression, or at least an important down-regulation, in mutant testes.

To determine the testicular localization of these lncRNA, *in situ* hybridization (*IS*H) on adult WT testes sections was performed (Supplementary Figure 7) and the results summarized (Figure 3B, D, F). These lncRNAs were expressed in spermatocytes and the most intense probe labeling was observed at the pachytene stage. These results were confirmed by data on the ReproGenomics viewer for *4930463O16Rik* and *Gm21269* (https://rgv.genouest.org/) (Supplementary Figures 3B and 4B) (Darde et al., 2015, 2019).

To refine the subcellular localization of these transcripts in adult mouse testes, we paired *IS*H experiments and the IF staining of DDX4 protein (or Mvh, Mouse Vasa homolog). DDX4 is a germ cell cytoplasmic marker of germ cells, especially in the testes (Toyooka et al., 2000). Our results showed that the three lncRNAs observed displayed different intensities of expression depending on seminiferous epithelium stages. *4930463O16Rik* was expressed in the nucleus of spermatocytes with diffuse fluorescence, surrounded by cytoplasmic DDX4 labelling from the zygotene to the diplotene stages (Figure 4A-B-C). At the same spermatocyte stages (zygotene to diplotene), a diffuse labelling of *Gm21269*, similar to that of *4930463O16Rik*, was observed but with the addition of dot-shaped labelling that co-localized with DDX4 fluorescence (Figure 4D, E, F). *Gm21269* was therefore localized in the cytoplasm and nuclei of spermatocytes during meiosis. *4921513H07Rik* appeared to be cytoplasmic, with fluorescent red dots (*IS*H) surrounding the nuclei, and located in close proximity to DDX4 (IF) labelling (Figure 4G, H, I). At other stages, identified by DDX4 staining, *IS*H labelling of these three lncRNA revealed single dots in a few spermatogonia and in round spermatids. The same experiment was then repeated: *IS*H was followed by IF staining of γH2Ax to highlight the sex body in spermatocytes (Supplementary Figure 8). No co-localization between the sex body and the three lncRNA was revealed.

**Figure 4:**
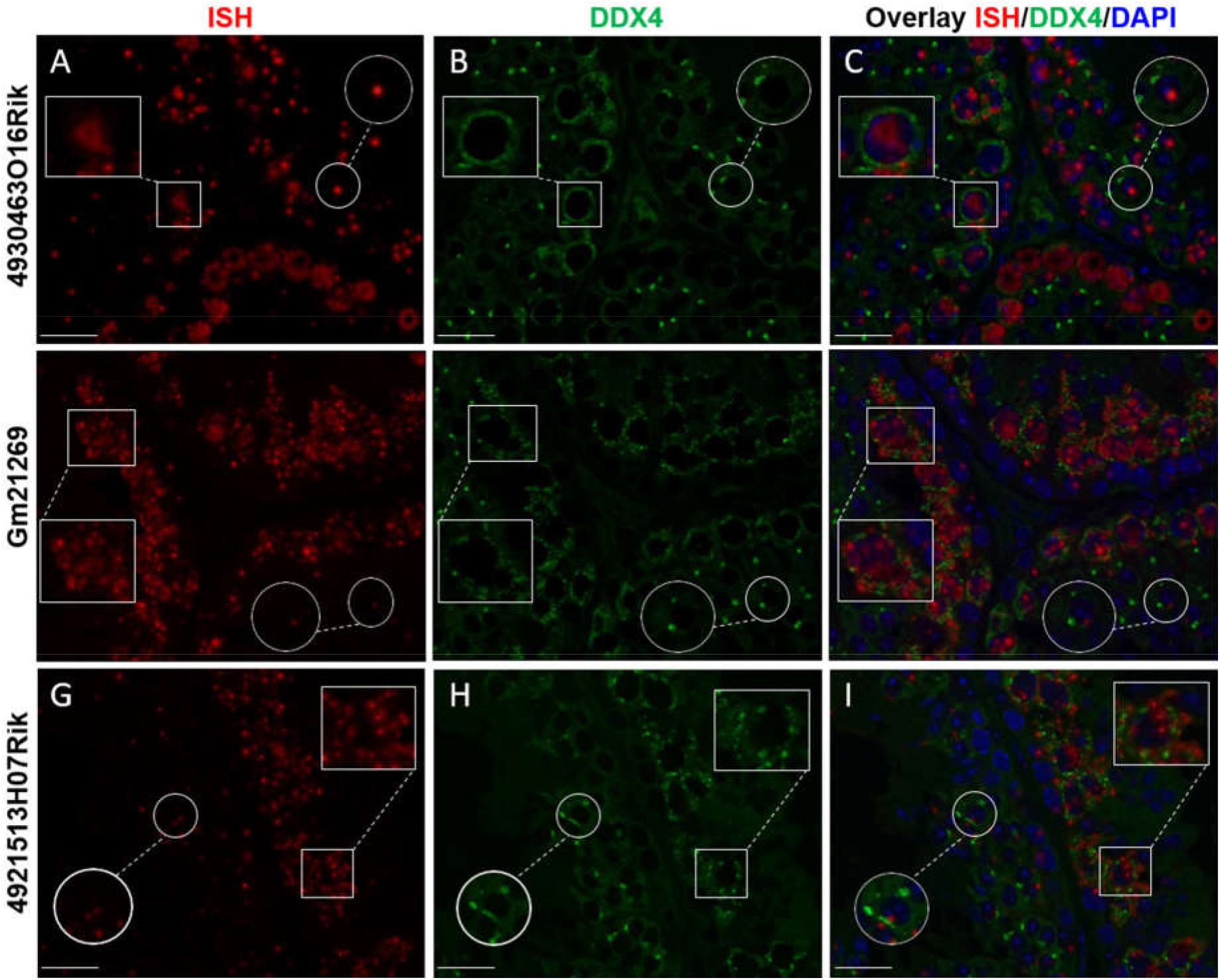
lncRNA cellular localizations on WT two month-old mouse testes. *In situ* hybridization using (A) *4930463O16Rik*, (D) *Gm21269* and (G) *4921513H07Rik* probes (red). (B-E-H) Immunofluorescence staining with DDX4 antibody was achieved at the same stage of seminiferous epithelium to identify male germ cells (green). (C-F-I) DAPI (blue), visualizing nuclear chromosomes, was merged with *IS*H (green) and IF (red) signals. Zooms in white squares show spermatocytes during the first meiotic division (zygotene to diplotene stages). Zooms in circles show spermatid cells with one spot of DDX4 staining per cell. Scale bar = 20 μm.

Taken together, these results indicate that these spermatocyte-specific lncRNAs had different subcellular localizations in spermatocytes, suggesting functions in these male germ cells.

### Generation of *4930463O16Rik*-deleted mice

In order to evaluate a potential role in spermatogenesis for one of these lncRNAs, *4930463O16Rik* (the nuclear expressed gene), it was decided to suppress its expression in a mouse knockout model. The *4930463O16Rik* gene (Chr10: 84,488,293-84,497,435 - GRCm38:CM001003.2) is described in public databases as consisting of four exons spanning approximately 10 kb in an intergenic locus on mouse chromosome 10. Using PCR and sequencing, we confirmed this arrangement (data not shown). In order to understand the role of *4930463O16Rik*, a new mouse line depleted of this lncRNA was created using CRISPR/Cas9 technology (Figure 5A). Briefly, multiple single guide RNAs (sgRNAs) were chosen, two sgRNAs in 5’ of exon 1 and two sgRNAs in 3’ of exon 4, so as to target the entire length of this gene (Figure 5A, C) and enhance the efficiency of gene deletion in the mouse (Han et al., 2014). Mice experiencing disruption of the target site were identified after the Sanger sequencing of PCR amplification of the genomic region surrounding the deleted locus (Figure 5D). *4930463O16Rik*^+/−^ mice were fertile and grew normally. Male and female *4930463O16Rik*^+/−^ animals were mated to obtain *4930463O16Rik*^−/−^ mice. Once the mouse line had been established, all pups were genotyped with a combination of primers (listed in Supplementary Table 4) (Figure 5B).

**Figure 5:**
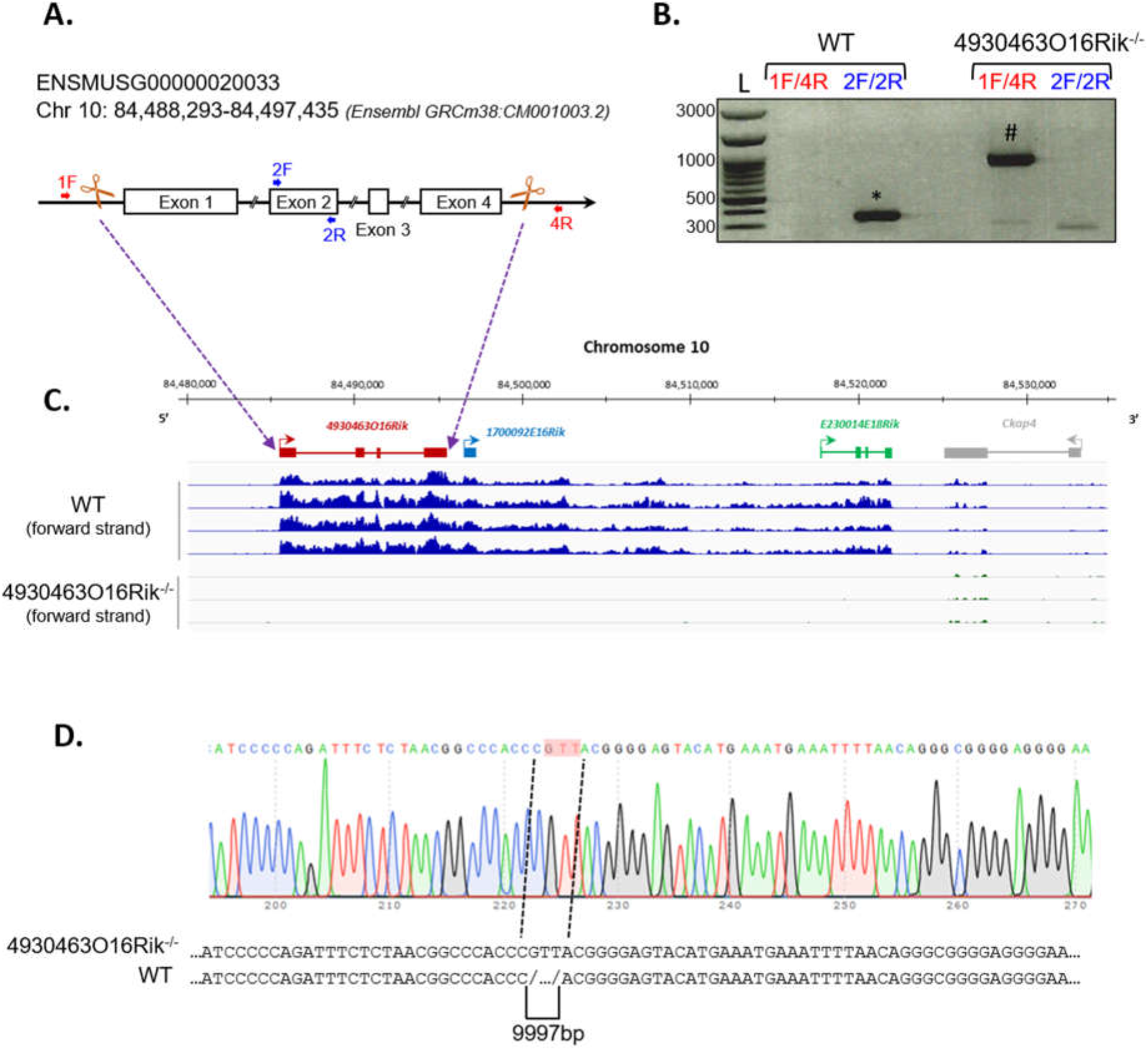
Deletion of the *4930463O16Rik* gene in the mouse. (A) Schematic design of CRISPR/Cas9 deletion of the *4930463O16Rik* gene with the suppression of 4 exons and 3 introns. The white boxes and lines represent exons and introns, respectively. (B) PCR genotyping on DNA of WT and *4930463O161Rik*^−/−^ mice. The primer pairs used 1F/4R (located in 5’ of exon 1 and in 3’ of exon 4 of the *4930463O16Rik* gene, respectively) or 2F/2R (located in the exon2 of *4930463O16Rik*) to determine the genotypes of the mice. Results showed the following amplicon sizes: (*) 352 bp with the 2F/2R primers in WT (no amplification in mutant mice); (#) 935 bp with the 1F/4R primers in *4930463O161Rik*^−/−^ mice (no amplification in WT mice under the PCR conditions used). (L) DNA ladder. (C) Transcription of the forward strand of chromosome 10 around the *4930463O16Rik* gene with RNA-seq coverage (BigWig format representation) in WT (top blue tracks) and *4930463O16Rik*^−/−^ (bottom tracks) mouse P18 testes. A continuous (WT) or very low transcription (*4930463O16Rik^−/−^*) was observed from *4930463O16Rik* to *E230014E18Rik* genes. (D) Electrophoregram of *4930463O16Rik^−/−^* mouse genomic DNA showing 9997 bp deletion and the insertion of 3 nucleotides (GTT, highlighted in pink).

### The absence of *4930463O16Rik* does not affect mouse fertility

Fertility was then investigated in *4930463O16Rik*-deficient mice. Eight-week-old male and female *4930463O16Rik^−/−^* mice were mated, and both sexes were fertile. Their litter sizes (7.5 ± 2.10 pups per litter, n=28) were similar to those of their WT counterparts (6.9 ± 2.12 pups per litter, n=20). There were no significant differences in terms of testicular size, testis morphology and histology and cauda and caput epididymis between WT and *4930463O16Rik^−/−^* adult mice (Figure 6A, B). In addition, the different stages of seminiferous tubules divided into seven groups were quantified between *4930463O16Rik^−/−^* and WT adult mice. No significant differences were observed between the two genotypes (Figure 6C). These results therefore demonstrated that *4930463O16Rik* is not required for mouse fertility.

**Figure 6:**
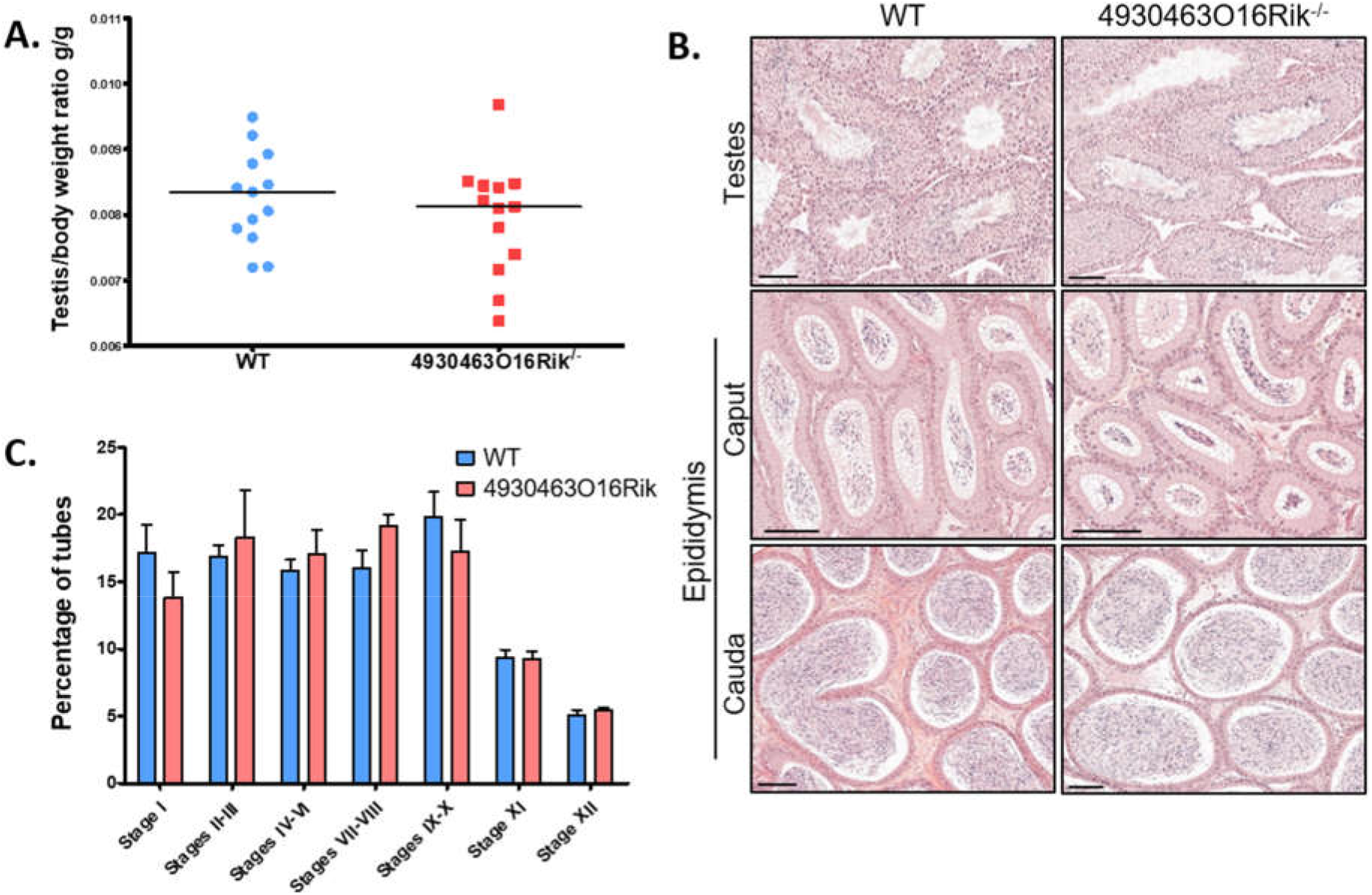
Study of *4930463O16Rik^−/−^* testicular phenotype. (A) Testis/body weight ratio of 8 week-old mice. No significant difference was observed in the two mouse lines. Median lines are in black. (B) Hematoxylin and eosin (HE) staining of testis and epididymis sections from WT and *4930463O16Rik^−/−^* 8 week-old mice. Scale bar=50 μm. Spermatozoa were visible in the lumen of the testes and epididymis of WT and *4930463O16Rik^−/−^* mice. (C) Quantification of the different seminiferous epithelium stages in WT and *4930463O16Rik^−/−^* 8 week-old mice. No significant difference was found between WT and *4930463O16Rik^−/−^* mice.

### *4930463O16Rik^−/−^* mice present modified sperm parameters

The sperm parameters of 8-week-old *4930463O16Rik*-deficient testes were compared to WT testes of the same age. Sperm concentrations obtained from the epididymis of *4930463O16Rik^−/−^* mice were significantly reduced by 57.2% compared to WT (Figure 7A) despite an unmodified testis/body weight ratio (Figure 6A). Motility parameters such as the percentage of motile spermatozoa, the motile mean expressed as beat cross frequency (bcf) and progressive spermatozoa were significantly higher in *4930463O16Rik^−/−^* mice compared to WT (Figure 7B, C, D). From a morphological point of view, however, two parameters were significantly modified in the testes of mutant mice: the distal mid-piece reflex (DMR), a defect developing in the epididymis and indicative of a sperm tail abnormality (Johnson, 1997) and the percentage of spermatozoa with coiled tail (Figure 7E, F). In addition, two kinetic parameters were also significantly reduced in mutant sperm: the motile mean vsl, related to the progressive velocity in a straight line and the average path velocity, or vap (Figure 7G, H).

**Figure 7:**
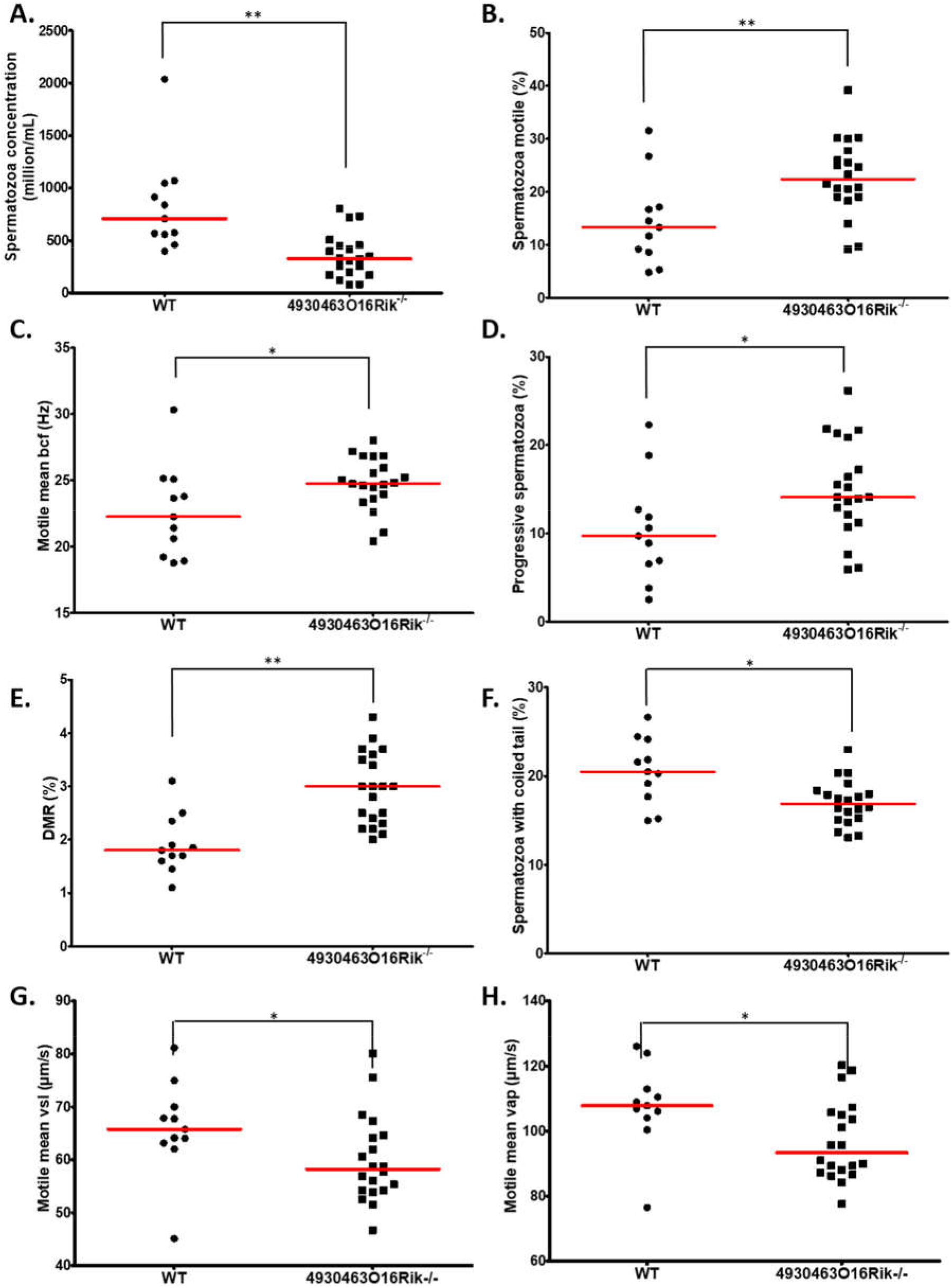
Evaluation of sperm parameters. Comparison of sperm-specific parameters from WT (circle, n=11) and *4930463O16Rik^−/−^* (square, n=20) mice. Significantly affected sperm parameters were (A) spermatozoa concentration (10^6^/mL), (B) spermatozoa motility (%), (C) motile mean bcf (beat cross frequency), (D) progressive spermatozoa (%), (E) DMR (distal midpiece reflex, abnormality of the sperm tail (%), (F) spermatozoa with coiled tail (%), (G) motile mean VSL (μm/s) and (H) VAP (μm/s). Statistical analyses were performed using the non-parametric Kruskal-Wallis test. * = p-val<0.05, ** = p-val<0.01.

These results obtained using computer-aided sperm analysis (CASA) thus showed that several sperm parameters; namely concentration, motility, morphology and kinetics were impacted in *4930463O16Rik* lncRNA-deficient mice. Some of them might negatively impact fertility, such as the sperm concentration, the DMR, the percentage of coiled tail and the motile mean vsl, while others would tend to suggest increased fertility, such as the motile mean percentage and bcf, and the progressive spermatozoa. These observations might have explained the normal fertility of *4930463O16Rik* lncRNA-deficient mice.

### Normal male fertility despite *4930463O16Rik^−/−^* mouse testis transcriptome modified

Transcriptomic RNA-seq analyses were performed in WT and *4930463O16Rik^−/−^* mouse testes at two developmental stages; i.e. at P16 and 18, as in the *Topaz1^−/−^* mouse line.

At P16, seven genes were differentially expressed (adjusted p-value<0.05; absolute Log2FC>1), including *4930463O16Rik, 1700092E16Rik* and *E230014E18Rik* (Supplementary Table 5). These latter two Riken cDNAs are in fact situated within the 3’ transcribed RNA of *4930463O16Rik* (positioned in Figure 5C) and correspond to a unique locus. The transcriptional activity of this new locus stops towards the 3’ end of the *cKap4* gene (cytoskeleton-associated protein 4 or Climp-63). This gene was down-regulated 1.7-fold in both knock-out lines (*Topaz1* and *4930463O16Rik*), which could suggest a newly discovered positive regulatory role for this lncRNA on the *cKap4* gene.

At P18, 258 genes were differentially expressed (199 down-regulated and 59 up-regulated using the same statistical parameters; Supplementary Table 5). Among them, 206 were protein-coding genes accounting for 79.8% of DEGs (Figure 8A). Thus, P18 DEGs highlighted a direct or indirect relationship between the loss of the *4930463O16Rik* lncRNA and protein-coding genes. In addition, loss of this lncRNA also resulted in the deregulation of 37 (14.3%) other lncRNAs.

**Figure 8:**
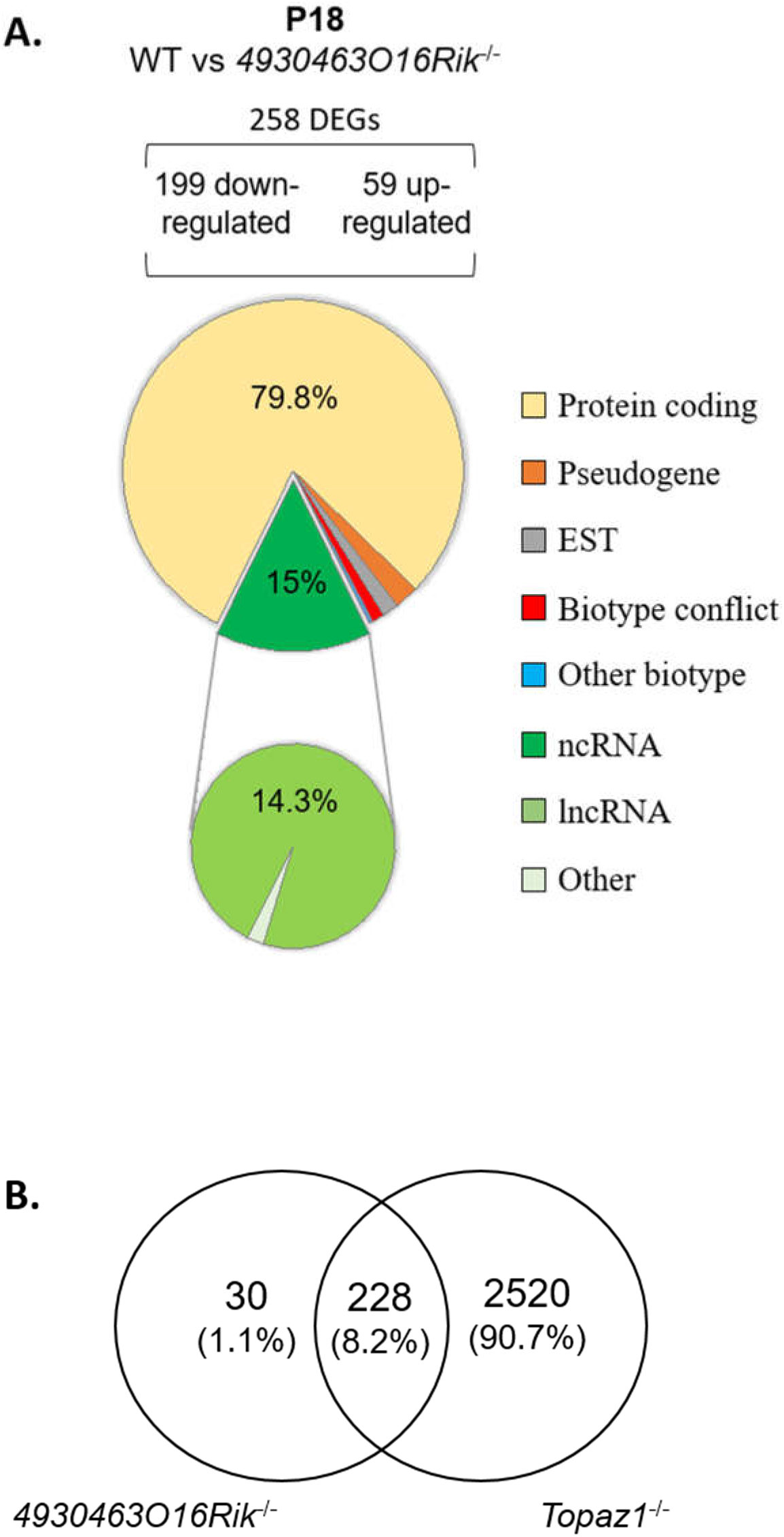
Deregulated genes from *4930463O16Rik*^−/−^ mouse testes. (A) Biotype of differentially expressed genes at P18 in *4930463O16Rik^−/−^* testes. Most of the deregulated genes are coding protein genes (adjusted p<0.05 and down-regulated FC<2 (log2FC<-1) or up-regulated FC>2 (log2FC>1). (B) Venn diagram showing the overlap of differentially expressed genes between *4930463O16Rik^−/−^* and *Topaz1^−/−^* mouse testes at 18 d*pp*.

The validation of several DEGs was performed by RT-qPCR using WT and *4930463O16Rik^−/−^* testicular RNAs at both developmental stages (P16 and P18) (Supplementary Figure 9). The qPCR results for the genes thus tested were consistent with those of RNA-seq.

The *4930463O16Rik^−/−^* DEGs were also analyzed using the DAVID database (Supplementary Table 6). At P18, six functional clusters had an enrichment score >1.3 (Huang et al., 2009a). As for *Topaz1^−/−^* mouse testes, they included the following GO terms: (*i*) for cellular components: cilium movement, ciliary part, axoneme, and (*ii*) for biological processes: microtubule-based process, regulation of cilium movement, spermatogenesis, male gamete generation, spermatid development and differentiation (Supplementary Table 6). An analysis that discriminated up-regulated from down-regulated genes only increased the value of the enrichment scores for the latter. The absence of TOPAZ1 protein or of *4930463O16Rik* lncRNA caused the same enrichment clusters in mutant testes despite different outcomes regarding the fertility of male mice. The other clusters from the DAVID analysis referred to the GO terms: cell surface, external side of membrane, defense or immune response and response to external stimulus. These clusters were only found in the DAVID analysis with up-regulated genes.

Therefore, the *4930463O16Rik* gene would appear to regulate genes related to spermatogenesis, microtubule or ciliary organization and the cytoskeleton in P18 testes. In the absence of this lncRNA, some genes involved in defense mechanisms or the immune response are also deregulated, suggesting stressful conditions. It should be noted that the majority (228/258 or 88%) of the DEGs from P18-*4930463O16Rik^−/−^* testes was in common with those deregulated in *Topaz1^−/−^* mice (Figure 8B). This led to similar results following ontological analyses of the DEGs of the two mutant lines. In the *Topaz1^−/−^* testes, these 228 genes could be a consequence of the down-regulation of *4930463O16Rik* lncRNA. On the other hand, these 228 DEGs alone do not explain meiotic arrest in the *Topaz1^−/−^* testes.

## Discussion

*Topaz1* was initially reported as a germ-cell specific factor (Baillet et al., 2011) essential for meiotic progression and male fertility in mice (Luangpraseuth-Prosper et al., 2015). The suppression of *Topaz1* led to an arrest of meiosis progression at the diplotene-metaphase I transition associated with germ cell apoptosis. Moreover, an initial transcriptomic approach, based on DNA microarrays, enabled the observation of a large but not exhaustive repertoire of deregulated transcripts. Using this technology, 10% of differentially-expressed probes were lncRNAs and presented deregulated expression in P20 *Topaz1^−/−^* testes compared to WT.

During this study, we were able to show that the effects of the*Topaz1* gene being absent were visible on the mouse testicular transcriptome as early as 16 days *post-partum*, i.e. before the first meiotic division and the production of haploid germ cells. These effects were amplified at 18 days *post-partum*, just before or at the very start of the first haploid germ cells appearing. The molecular pathways involved in the suppression of TOPAZ1 form part of spermatogenesis and the establishment of the cell cytoskeleton. At these two stages, P16 and P18, about a quarter of the deregulated genes in testes were ncRNAs (mainly lncRNAs) some of which displayed almost no expression in *Topaz1^−/−^* testes. Suppressing one of them did not prevent the production of haploid spermatids and spermatozoa, but halved the murine sperm concentration. Furthermore, by deleting ~10 kb corresponding to this *4930463O16Rik* lincRNA, we showed that the transcriptional extinction was even longer, encompassing ~35 kb in total and two other genes (1700092E16Rik (unknown gene type according to Ensembl) and the lincRNA E230014E18Rik). Indeed, our transcriptional data suggested that these three annotated loci belong to a unique gene. Transcription of this lincRNA ended near the 3’ region of the *cKap4* gene, known to be associated with the cytoskeleton (Vedrenne and Hauri, 2006). Remarkably, cKap4 expression was down-regulated 1.7 fold in both types of knockout mouse (*Topaz1^−/−^* and *4930463O16Rik^−/−^*), suggesting a previously unknown and positive regulatory role of *4930463O16Rik* on *cKap4*.

### *Topaz1* ablation leads to chromosome misalignments at pro-metaphase I

Meiosis and its two cell divisions are well-orchestrated sequences of events controlled by different genes. Although these divisions have many similarities between males and females, meiosis is also sex-dimorphic. This particularly concerns timing, synchronization, the number of haploid gametes produced and the periods of meiotic arrest (reviewed by (Handel and Eppig, 1998)). In females, meiosis is initiated during fetal life and oocyte development is arrested at the end of prophase I; they remain in this arrested state until the onset of ovulatory cycles around puberty. The first division of meiosis then resumes and leads to the release of a first polar globule with the secondary oocyte. At metaphase II, the oocyte is blocked again. Release of the second polar globule leading to the formation of the female gamete only occurs at fertilization (Handel and Eppig, 1998). In males, meiosis is a continuous process during the post-natal period, just before puberty, and results in the formation of four male gametes from one spermatocyte. Despite these sex-dimorphic differences, the first reductional division of meiosis is highly conserved between species and between sexes in terms of morphology and genetic regulation. It has been hypothesized that the mechanisms regulating and controlling prophase I during mammalian meiosis, frequently named “checkpoints”, are more stringent in males than in females. This has been demonstrated during the past 25 years by the use of a large number of mutant mouse models, mainly gene knockout mice (Morelli and Cohen, 2005; Handel and Schimenti, 2010; Su et al., 2011).

A major checkpoint in males is the synaptic checkpoint that controls zygotene-pachytene transition, highlighted in male mice lacking *Sycp3*, *Dmc1*, *Spo11*, *mei1*, *Msh4*-*5* or *OvoL1* genes (Pittman et al., 1998; Edelmann et al., 1999; Baudat et al., 2000; Kneitz et al., 2000; Romanienko and Camerini-Otero, 2000; Yuan et al., 2000; Li et al., 2005; Reinholdt and Schimenti, 2005). In mutant females, this synaptic checkpoint is less stringent. Indeed, female meiotic arrest may occur later, starting from the diplotene, as seen in *Dmc1*, *mei1*, *Msh4*-*5* knockout mice during the dictyate-resting phase of the oocyte, evidenced in *Spo11*. These female mice may even be fertile, as seen in *OvoL1* knockout mouse (Li et al., 2005). Other gene suppressions have highlighted a second meiotic checkpoint of metaphase I in males (such as those of *Mlh1-3*, or cyclin A1) due to a misalignment of chromosomes on the spindle (Liu et al., 1998; Eaker et al., 2002; Lipkin et al., 2002). Male mice devoid of the *Topaz1* gene may match these latter models. Indeed, *Topaz1^−/−^* spermatocytes do not progress to metaphase I and the chromosomes are not properly aligned on the metaphase plate. *Topaz1^−/−^* females are however fertile.

### TOPAZ1 seems to be involved in the shape, structure and movements of cells

Absence of the *Topaz1* gene disturbs the transcriptome of murine testes as early as 16 days postnatal. Of the 205 DEGs at P16, 85 were specific to this stage of development compared to P18 (Figure 1A), such as *Ptgs1* (*Cox1*) a marker of peritubular cells (Rey-Ares et al., 2018), and *Krt18* a marker of Sertoli cell maturation (Tarulli et al., 2006). Moreover, different genes involved in the TGFβ pathway were also P16-DEGs, such as *Bmpr1b*, *Amh* and *Fstl3*. The latter, for example, was demonstrated to reduce the number of Sertoli cells in mouse testes and to limit their size (Oldknow et al., 2013). *Ptgds* (L-*Pgds*), which plays a role in the PGD2 molecular pathway during mammalian testicular organogenesis, is also deregulated (Moniot et al., 2009). All these genes, specifically deregulated at P16 due to the absence of *Topaz1,* thus appeared to participate in regulating cell communication. At P18, these genes were no longer differential but they could be replaced by genes belonging to the same gene families, such as the cadherin or keratin families.

Many of the 205 DEGs at P16 were involved in defense response pathways. For example, *Ifit3* and *Gbp3* are immune response genes in spermatocyte-derived GC-2spd(ts) cells (Kurihara et al., 2017). Two days later, at P18, just before the prophase I-metaphase 1 transition, ten times more genes were deregulated. Among the 120 DEGs common to P16 and P18, there was at least one gene that might be involved in meiosis such as *Aym1*, an activator of yeast meiotic promoters 1. The absence of *Topaz1* led to a lack of the testicular expression of *Aym1*. This gene is germ cell-specific (Malcov et al., 2004). In male mice, *Aym1* is expressed from 10 d*pp* in early meiotic spermatocytes. The small murine AYM1 protein (44 amino acids) is immunolocalized in the nucleus of primary spermatocytes, mainly late pachytene and diplotene, suggesting a nuclear role for AYM1 in germ cells during the first meiotic division(Malcov et al., 2004).

At P18, the testicular transcriptome of *Topaz1^−/−^* mice was largely disturbed when compared to WT animals, and most DEGs were down-regulated (Figure 1F), suggesting that TOPAZ1 promotes gene expression in normal mice. As TOPAZ1 is predicted to be an RNA-binding protein, it is tempting to speculate that its absence disorganized ribonucleic-protein complexes, including their instabilities and degradation. This could partly explain why 90% of DEGs were down-regulated at P18 and included a large proportion of lincRNAs. These down-regulated genes at P18 concerned microtubule-based movement and microtubule-based processes, and cellular components relative to motile cilium, ciliary part, sperm flagellum and axoneme. In addition, DAVID analysis revealed GO terms such as centriole, microtubule and spermatogenesis. All these terms relate to elements of the cytoskeleton that are indispensable for mitotic and/or meiotic divisions, motility and differentiation and are also widely involved in spermiogenesis, as might be expected with this latter GO term because most DEGS are testis-specific. The centriole is a widely conserved organelle in most organisms. A pair of centrioles is located at the heart of the centrosome, and the whole is grouped together as the main microtubule-organizing center (MTOC). During our study, staining of the meiotic spindle and centrosomes revealed a disturbance of these pathways (Figure 2 and Supplementary Figure 2). Such abnormal metaphase-like chromosomes were arranged in rosettes rather than being neatly aligned at the cell equator, and hemispindles centered in the spermatocytes had previously been observed. For example, aberrant prometaphase-like cells were observed in *Mlh1*- or *Meioc*-deficient testes (*Meioc* is down-regulated 1.51-fold in P18 in *Topaz1^−/−^* testes) (Eaker et al., 2002; Abby et al., 2016). These mutant mice have been described as displaying an arrest of male meiosis, and testes devoid of haploid germ cells leading to male sterility like mice lacking the *Topaz1* gene. In *Topaz1^−/−^* testes, Mlh1 is not a DEG.

During spermatogenesis, the dysregulation of centrosome proteins may affect meiotic division and genome stability. The centriole proteins CEP126, CEP128, CEP63 were down-regulated (FC from 2.1 to 2.7 compared to WT) at P18 in *Topaz1^−/−^* testes. CEP126 is localized with γ-tubulin on the centriole during the mitosis of hTERT-RPE-1 (human telomerase-immortalized retinal pigmented epithelial cells) (Bonavita et al., 2014) but has never been studied in germ cells during meiosis. CEP128 was localized to the mother centriole and required to regulate ciliary signaling in zebrafish (Mönnich et al., 2018). *Cep128* deletion decreased the stability of centriolar microtubules in F9 cells (epithelial cells from testicular teratoma of mouse embryo) (Kashihara et al., 2019). Centriole separation normally occurs at the end of prophase I or in early metaphase I, and CEP63 is associated with the mother centrioles. The mouse model devoid of *Cep63* leads to male infertility (Marjanović et al., 2015), and in spermatocytes from these mice, the centriole duplication was impaired. Finally, our ontology analysis of *Topaz1^−/−^* P18-DEGs revealed significant enrichment scores for the several clusters relative to the final structure of spermatozoa such as tetratricopeptide repeat (TPR) and dynein heavy chain (DNAH1) clusters. Dynein chains are macromolecular complexes connecting central or doublet pairs of microtubules together to form the flagellar axoneme, the motility apparatus of spermatozoa (ref in (Miyata et al., 2020a)). Dynein proteins have also been identified as being involved in the microtubule-based intracellular transport of vesicles, and in both mitosis and meiosis (Mountain and Compton, 2000).

The TPR or PPR (pentatricopeptide repeat) domains consist of several 34 or 36 amino acid repeats that make up αα-hairpin repeat units, respectively (D’Andrea and Regan, 2003). The functions of TPR or PPR proteins were firstly documented in plants and are involved in RNA editing (D’Andrea and Regan, 2003; Schmitz-Linneweber and Small, 2008). In the mouse, *Cfap70*, a tetratricopeptide repeat-containing gene, was shown to be expressed in the testes (Shamoto et al., 2018), or as *Spag1* in late-pachytene spermatocytes or round spermatids (Takaishi and Huh, 1999). Moreover, *Ttc21a* knockout mice have displayed sperm structural defects of the flagella and the connecting piece. In humans, *Ttc21a* has been associated with asthenoteratospermia in the Chinese population (Liu et al., 2019).

Numerous components of the intraflagellar transport (IFT) complex contain TPR. Several genes coding for such tetratricopeptide repeat-containing proteins are down-regulated in P18 testes devoid of *Topaz1,* such as *Cfap70*, *Spag1*, *Tct21a* and *Ift140*. Based on TPRpred (Karpenahalli et al., 2007) that predicted TPR- or PPR-containing proteins, the TOPAZ1 protein was predicted to contain such domains; seven in mice (p-val= 7.5E-08, probability of being PPR= 46.80%) and ten in humans (p-val= 3.4E-09, probability of being PPR = 88.76%).

A recent study of single cell-RNA-seq from all types of homogeneous spermatogenetic cells identified clusters of cells at similar developmental stages (Chen et al., 2018a). This study shown that most of the genes involved in spermiogenesis start being expressed from the early pachytene stage. This is consistent with our RNA-seq results. Taken together, these data indicate that the absence of *Topaz1* down-regulated a significant number of cytoskeleton-related genes, leading to a defect in formation of the meiotic spindle and to a deficient duplication and/or migration of centrosomes as early as 18 days post-natal. *Topaz1* could lead to impaired chromosome dynamics via the activation of cytoskeleton genes, thus revealing the essential role of the centrosome in promoting division and then fertility. TOPAZ1 may act via its TPR domains.

### *Topaz1* ablation deregulates a high proportion of lncRNAs

DEGs between *Topaz1^−/−^* and WT mouse testes also revealed a high proportion of deregulated lncRNAs. We showed that three lincRNAs, whose expression was almost abolished as early as P16 in *Topaz1-deficient* mouse testes, were testis- and germ cell-specific. We showed that these genes are expressed in spermatocytes and round spermatids, suggesting a role in spermatogenesis. Their functions are still unknown.

Several investigations have revealed that the teste allow the expression of many lncRNAs (Necsulea et al., 2014). In mammals, the testis is the organ with the highest transcription rate (Soumillon et al., 2013). However, during the long stage of prophase I, these levels of transcription are not consistent. Indeed, transcription is markedly reduced or even abolished in the entire nucleus of spermatocytes during the early stages of prophase I. This is accompanied in particular by the nuclear processes of DNA division, the pairing of homologous chromosomes and telomeric rearrangements (Bolcun-Filas and Schimenti, 2012; Baudat et al., 2013; Shibuya and Watanabe, 2014), and also by the appearance of MSCI (meiotic sex chromosome inactivation) markers (Page et al., 2012). These processes are supported by epigenetic changes such as histone modifications and the recruitment of specific histone variants (references in Page et al., 2012). Transcription then takes up an important role in late-pachytene to diplotene spermatocytes (Monesi, 1964). The aforementioned scRNA-seq study of individual spermatogenic cells showed that almost 80% of annotated autosomal lncRNAs were expressed in spermatogenetic cells, mainly in mid-pachytene- to metaphase I-spermatocytes but also in round spermatids (Chen et al., 2018a). The three lncRNAs investigated during our study (*4930463O16Rik, Gm21269* and *4921513H07Rik*) were also expressed at these developmental stages in mouse testes (Chen et al., 2018b; Li et al., 2021). In the latter study (Li et al., 2021), the authors identified certain male germline-associated lncRNAs as being potentially important to spermatogenesis *in vivo*, based on several computational and experimental data sets; these lncRNAs included *Gm21269* and *4921513H07Rik*. The localization of lncRNAs in cells may be indicative of their potential function (Chen, 2016). *4930463O16Rik* is expressed in the nucleus of spermatocytes. As mentioned above, *4930463O16Rik* may play a positive role in the expression of *cKap4* at the neighboring locus. Some nuclear lncRNA are involved in regulating transcription with a *cis*-regulatory role, such as *Malat1* or *Air* (Sleutels et al., 2002; Zhang et al., 2012) on a nearby gene. Other nuclear lncRNAs act in *trans* and regulate gene transcription at another locus, such as *HOTAIR* (Chu et al., 2011). In addition, some cytoplasmic lncRNA have been shown to play a role in miRNA competition, acting as miRNA sponges or decoys (such as *linc-MD1* in human myoblasts (Cesana et al., 2011)). *Gm21269* is localized in the cytoplasm and nuclei of spermatocytes during meiosis. Both cytoplasmic and nuclear lncRNAs may act as a molecular scaffold for the assembly of functional protein complexes, such as *HOTAIR* or *Dali* (Tsai et al., 2010; Chalei et al., 2014), regulating protein localization and/or direct protein degradation, or acting as an miRNA precursor (Cai and Cullen, 2007). Finally, multiple other roles can be observed for lncRNAs. For example, the *Dali* lincRNA locally regulates its neighboring *Pou3f3* gene, acts as a molecular scaffold for POU3F3 protein and interacts with DNMT1 in regulating the DNA methylation status of CpG island-associated promoters in *trans* during neural differentiation (Chalei et al., 2014).

### The deletion of one lncRNA alters sperm parameters without affecting fertility

To decipher the biological function of an lncRNA affected by *Topaz1* invalidation, a mouse model devoid of *4930463O16Rik* was produced, with the same genetic background as *Topaz1^−/−^* mice. This knockout mouse model did not exhibit meiosis disruption and the fertility of these mutant mice remained intact under standard laboratory conditions. Using a similar approach, *Sox30* is a testis-specific factor that is essential to obtain haploid germ cells during spermatogenesis (Bai et al., 2018). SOX30 regulates *Dnajb8* expression, but the deletion of *Dnajb8* is not essential for spermatogenesis and male fertility (Wang et al., 2020).

Several mutant mice deprived of testis-specific genes proved to be fertile, although no role has been established for these genes during spermatogenesis. This was noted in particular for the *Flacc1*, *Trim69*, *Tex55*, *4930524B15Rik* genes (Chotiner et al., 2020; He et al., 2020; Khan et al., 2020; Jamin et al., 2021) or for highly testis-enriched genes such as *Kdm4d*, *Tex37*, *Ccdc73* or *Prss55* (Iwamori et al., 2011; Khan et al., 2018). Some of them were down-regulated genes in *Topaz1^−/−^* or *4930463O16Rik^−/−^* testes (*Trim69* in *Topaz1^−/−^* FC = 3.99 and in *4930463O16Rik^−/−^* FC = 2.27; *Kdm4d* in *Topaz1^−/−^* FC = 2.70 and in *4930463O16Rik^−/−^* FC = 1.83; *Ccdc73* in *Topaz1^−/−^* FC = 1.46). Some laboratories have recently also generated several dozen testis-enriched knockout mouse lines using the CRISPR/Cas9 system and shown that all these genes are individually dispensable in terms of male fertility in mice (Miyata et al., 2016; Lu et al., 2019).

The abundant expression of lncRNAs during spermatogenesis has also prompted other laboratories to produce knockout mouse models of testis-specific lncRNAs. This was the case for *1700121C10Rik* or *lncRNA5512* lncRNAs where mutant mice were also fertile without variations in their sperm parameters (Li et al., 2020; Zhu et al., 2020). One working hypothesis might be that some lncRNAs may regulate subsets of functional spermatogenetic-gene expression, in line with their nuclear localization, by binding to their regulatory genomic region.

Nevertheless, in our *4930463O16Rik*-knockout mouse model, several sperm parameters were altered, including reduction in epididymal sperm concentrations (by more than half) and sperm motility. In *Tslrn1* knockout mice (testis-specific long non-coding RNA 1) the males were fertile and displayed significantly lower sperm levels (−20%) but no reduction in litter size, or major defects in testis histology or variations in sperm motility (Wichman et al., 2017). In *Kif9*-mutant male mice, no testes abnormalities were found (Miyata et al., 2020b). They were sub-fertile due to impaired sperm motility: the VSL and VAP velocity parameters were reduced, as in *4930463O16Rik* knockout mice. The authors concluded that *Kif9* mutant mice were still fertile and this was probably due to variations in the motility of individual spermatozoa; those with good motility could still fertilize oocytes. The same conclusion may apply to *4930463O16Rik^−/−^* mice.

The suppression of a gene – in this case *4930463O16Rik* lincRNA – whose expression is markedly down-regulated in the testes of sterile *Topaz1^−/−^* mice (FC = 40), has no effect on spermatogenesis. Our data suggest that the expression of *4930463O16Rik* is not essential for meiotic division but adds to the terminal differentiation of male germ cells.

Various genes, either testis-specific or highly expressed in the testes, exert no effect on reproduction when deleted independently (Miyata et al., 2016; Li et al., 2020). Given the large number of lncRNAs expressed in meiotic testes, one explanation may be that the function of *4930463O16Rik* is partly redundant with that of other testicular lncRNAs.

However, outside the laboratory, in wild reproductive life, one might imagine that biological functions may differ under more natural conditions due to stress and reproductive competition. This has been shown in particular for *Pkdrej*-deficent male mice which are fertile, whereas the *Pkdrej* gene (polycystin family receptor for egg jelly), is important to postcopulatory reproductive selection (Sutton et al., 2008; Miyata et al., 2016).

The absence of a specific anti-TOPAZ1 antibody did not enable us to further advance in our understanding of its function during murine spermatogenesis. The creation of a Flag-tagged *Topaz1* knockin mouse model will allow us to gain further insights, and Rip-seq experiments will enable the determination of RNA-TOPAZ1 complexes during spermatogenesis.

In summary, *Topaz1* is a gene that is essential for fertility in male mice. Its absence leads to meiotic arrest before the first division; germ cells display a centered monopolar spindle and a misarrangement of chromosomes. In addition, *Topaz1* stabilizes the expression of many lncRNAs. The suppression of one of them is not essential to mouse fertility but it is necessary during the terminal differentiation of male germ cells to achieve optimal function.

## Materials and methods

### Ethics statement

All animal experiments were performed in strict accordance with the guidelines of the Code for Methods and Welfare Considerations in Behavioral Research with Animals (Directive 2016/63/UE). All experiments were approved by the INRAE Ethical Committee for Animal Experimentation covering Jouy-en-Josas (COMETHEA, no. 18-12) and authorized by the French Ministry for Higher Education, Research and Innovation (No. 815-2015073014516635).

### Mice

The generation and preliminary analysis of *Topaz1*-null transgenic mouse line has been described previously (Luangpraseuth-Prosper *et al*. 2015).

Generation of the *4630493O16Rik*-null transgenic mouse line was achieved using CrispR-Cas9 genome editing technology. The RNA mix comprised an mRNA encoding for SpCas9-HF1 nuclease and the four sgRNA (Supplementary Table 4) targeting the *4930463o16Rik* gene (NC_000076: 84324157-84333540). These sgRNAs were chosen according to CRISPOR software (http://crispor.tefor.net/) in order to remove the four exons and introns of the *4930463o16Rik* gene. Cas9-encoding mRNA and the four sgRNAs were injected at a rate of 100 ng/μL each into one cell fertilized C57Bl/6N mouse eggs (Henao-Mejia et al., 2016).

The surviving injected eggs were transferred into pseudo-pregnant recipient mice. Tail-DNA analysis of the resulting live pups was performed using PCR with genotyping oligonucleotides (Supplementary Table 4) and the Takara Ex Taq^®^ DNA Polymerase kit. The PCR conditions were 94 °C 30s, 60 °C 30s and 72 °C 30s, with 35 amplification cycles.

Two transgenic founder mice were then crossed with wild-type C57Bl/6N mice to establish transgenic lines.

F1 heterozygote mice were crossed together in each line to obtain F2 homozygote mice, thus establishing the 4630493O16Rik^−/−^ mouse lines. Both mouse lines were fertile and the number of pups was equivalent, so we worked with one mouse line.

All mice were fed *ad libitum* and were housed at a temperature of 25°C under a 12h/12h light/dark cycle at the UE0907 unit (INRAE, Jouy-en-Josas, France). The animals were placed in an enriched environment in order to improve their receptiveness while respecting the 3R. All mice were then sacrificed by cervical dislocation. Tissues at different developmental stages were dissected and fixed as indicated below, or flash frozen immediately in liquid nitrogen before storage at −80°C. The frozen tissues were used for the molecular biology experiments described below.

### Histological and immunohistochemical analyses

For histological studies, fresh tissues from 8-week-old mice were fixed in 4% paraformaldehyde (Electron Microscopy Sciences reference 50-980-495) in phosphate buffer saline (PBS) at 4°C. After rinsing the tissues in PBS, they were stored in 70% ethanol at 4°C. Paraffin inclusions were then performed using a Citadel automat (Thermo Scientific Shandon Citadel 1000) according to a standard protocol. Tissues included in paraffin blocks were sectioned at 4μm and organized on Superfrost Plus Slides (reference J1800AMNZ). Once dry, the slides were stored at 4°C. On the day of the experiment, these slides of sectioned tissues were deparaffinized and rehydrated in successive baths of xylene and ethanol at room temperature. For histology, testes sections were stained with hematoxylin and eosin (HE) by the @Bridge platform (INRAE, Jouy-en-Josas) using an automatic Varistain Slide Stainer (Thermo Fisher Scientific). Periodic acid-Schiff staining (PAS) was used to determine seminiferous epithelium stages.

*In situ* hybridization experiments were performed using the RNAscope^®^ system (ACB, Bio-Techne SAS, Rennes, France). Briefly, probes (around 1000 nt long) for *Topaz1* (NM_001199736.1), *4930463o16Rik* (NR_108059.1), *Gm21269* (NR_102375.1) and *4921513H07Rik* (NR_153846.1) were designed by ACB and referenced with the catalog numbers 402321, 431411, 549421 and 549441, respectively. Negative and positive controls were ordered from ACD with *Bacillus subtilis* dihydrodipicolinate reductase (dapB) and *Homo sapiens* ubiquitin C (Hs-UBC), respectively. Hybridization was performed according to the manufacturer’s instructions using a labelling kit (RNAscope^®^ 2.5HD assay-brown). Brown labelling slides were counterstained according to a PAS staining protocol and then observed for visible signals. Hybridization was considered to be positive when at least one dot was observed in a cell. Stained sections were scanned using a 3DHISTECH panoramic scanner at the @Bridge platform (INRAE, Jouy-en-Josas) and analyzed with Case Viewer software (3DHISTECH). We also used the RNAscope^®^ 2.5HD assay-red kit in combination with immunofluorescence in order to achieve the simultaneous visualization of RNA and protein on the same slide. The *IS*H protocol was thus stopped by immersion in water before hematoxylin counterstaining. Instead, the slides were washed in PBS at room temperature. The Mouse on mouse (M.O.M.) kit (BMK-2202, Vector laboratories) was used and slides were incubated for one hour in Blocking Reagent, 5 minutes in Working solution and 2 hours with a primary antibody: DDX4 (ab13840, Abcam) or γH2AX(Ser139) (Merck), diluted at 1:200 in Blocking Reagent. Detection was ensured using secondary antibody conjugated to DyLight 488 (green, KPL). Diluted DAPI (1:1000 in PBS) was then applied to the slides for eight minutes. The slides were then mounted with Vectashield Hard Set Mounting Medium for fluorescence H-1400 and images were captured at the MIMA2 platform (https://www6.jouy.inrae.fr/mima2/, https://doi.org/10.15454/1.5572348210007727E12) using an inverted ZEISS AxioObserver Z1 microscope equipped with an ApoTome slider, a Colibri light source and Axiocam MRm camera. Images were analyzed using Axiovision software 4.8.2 (Carl Zeiss, Germany).

### Total RNA extraction and Quantitative RT-PCR (RT-qPCR)

Total RNAs from post-natal mouse testes or other organs were isolated using Trizol reagent. The RNAs were purified using the RNeasy Mini kit (Qiagen) following the manufacturer’s instructions and then DNAse-treated (Qiagen). The quantification of total RNAs was achieved with a Qbit^®^ Fluorometric Quantitation. Maxima First-Strand cDNA Synthesis Kit (Thermo Scientific) was used to reverse transcript RNA into cDNA. The Step One system with Fast SYBR− Green Master Mix (Applied Biosystems, ThermoFisher France) was used for qPCR, which was performed in duplicate for all tested genes and the results were normalized with qBase^+^ software (Biogazelle) (Hellemans et al., 2007). Gapdh, Ywahz and Mapk1 were used as the reference genes. For each experiment, median values were plotted using GraphPad Prism, and statistical analyses were performed with KrusKall-Wallis tests under R software (Rcmdr package (p-value<0.05)). The primer sequences used for RT-qPCR are shown in Supplementary Table 4.

### RNA-sequencing

Total RNA quality was verified on an Agilent 2100 Bioanalyser (Matriks, Norway) and samples with a RIN>9 were made available for RNA-sequencing. This work benefited from the facilities and expertise of the I2BC High-throughput Sequencing Platform (Gif-sur-Yvette, Université Paris-Saclay, France) for oriented library preparation (Illumina Truseq RNA Sample Preparation Kit) and sequencing (Paired-end 75 bp; NextSeq). More than 38 million 75 bp paired-end reads per sample were generated.

### Transcriptomic analysis

Sequence libraries were aligned with the Ensembl 95 genome using TopHat (Trapnell et al., 2009), and gene table counts were obtained by applying featureCounts to these alignments (Liao et al., 2014). Data normalization and single-gene level analyses of differential expression were performed using DESeq2 (Love et al., 2014). Some samples were sequenced several months apart. A batch effect was observed after computation of the hierarchical clustering of samples. In order to take this effect into account, we introduced the batch number into the DESeq2 model, as well as the study conditions. Differences were considered to be significant for Benjamini-Hochberg adjusted p-values <0.05, and absolute fold changes >2 (absolute Log2FC>1) (Benjamini and Hochberg, 1995). Raw RNA-seq data were deposited via the SRA Submission portal (https://submit.ncbi.nlm.nih.gov/subs/sra/), BioProject ID PRJNA698440.

### Biotype determination of DEGs

Data available on the NCBI, MGI (http://www.informatics.jax.org) and Ensembl (https://www.ensembl.org/) websites were used simultaneously to determine the DEG biotypes. For this purpose, information on the mouse genome was obtained by ftp from NCBI (ftp://ftp.ncbi.nih.gov/gene/DATA/GENE_INFO/Mammalia/Mus_musculus.gene_info.gz); the annotation BioMart file from Ensembl (http://www.ensembl.org/biomart/martview; Ensembl genes 95, Mouse genes GRCm28.p6) and feature types from MGI (http://www.informatics.jax.org/marker/; with the protein coding gene, non-coding RNA gene, unclassified gene and pseudogenic region). Only data corresponding to the DEGs were conserved. The files from these three databases were therefore cross-referenced to determine DEG biotypes. When the biotype of a gene differed between databases, the annotation was then listed as genes with a “biotype conflict”.

### Gene ontology enrichment

The mouse DEGS thus identified were analyzed through Gene Ontology (GO) and Kyoto Encyclopedia of Genes and Genomes (KEGG) pathway membership with Database performed using the DAVID Bioinformatic Database 6.8 (https://david.ncifcrf.gov/). These analyses and pathways were considered to be significant for a Benjamini-corrected enrichment p-value of less than 0.05. The Mouse Atlas Genome of differentially expressed genes extracted from this study was performed via the Enrichr website (https://maayanlab.cloud/Enrichr/).

### Sperm analysis

Evaluations of the concentrations and motility of sperm in WT and *4930463O16Rik^−/−^* 8-week-old mice were performed using the IVOS II Computer Assisted Sperm Analysis (CASA) system (Hamilton Thorne, Beverly, MA, USA). The two fresh cauda epididymes from each individual were removed and plunged into 200 μL TCF buffer (Tris, citrate and fructose buffer) where they were chopped up with small scissors. For sperm release, the samples were incubated for 10 minutes at 37°C. A 4 μl aliquot was placed in a standardized four-chamber Leja counting slide (Leja Products B.V., Nieuw-Vennep, Netherlands). Ten microscope fields were analyzed using the predetermined starting position within each chamber with an automated stage. Statistical analyses were performed using the mean of the 10 analyzed fields containing at least 300 cells. The IVOS settings chosen were those defined for mouse sperm cell analysis (by Hamilton Thorne). The principal parameters were fixed as follows: 45 frames were captured at 60 Hz; for cell detection, the camera considered a signal as a spermatozoon when the elongation percentage was between 70 (maximum) and 2 (minimum); the minimal brightness of the head at 186, and the minimum and maximum size of the head at 7 and 100 μm², respectively. The kinematic thresholds applied were: cell travel max at 10μm, progressive STR at 45%, progressive VAP at 45μm/s, slow VAP at 20μm/s, slow VSL at 30μm/s, static VAP at 4μm/S and static VSL at 1μm/s. The full settings used are listed in Supplementary Table 7. The CASA parameters thus recorded included the average path velocity (VAP in μm/s), straight line velocity (VSL in μm/s), curvilinear velocity (VCL in μm/s), amplitude of lateral head displacement (ALH in μm), motility (percentage), and sperm concentration (.10^6^/mL). Slow cells were recorded as static. Median and interquartile ranges were plotted with GraphPad. To compare the sperm parameters between WT and *4930463O16Rik^−/−^* mice, statistical analyses were performed using the Kruskal-Wallis non-parametric test.

## Acknowledgments

Our warmest thanks go to Jean-Luc Vilotte for allowing lncRNA deletion to take place in his laboratory, and also for proofreading this manuscript. We would like to thank the TACGENE facility at U1154-UMR7196, MNHN, Paris, and especially Anne De Cian and Jean-Paul Concordet, for the synthesis of gRNAs and Cas9 mRNA. We are grateful to all members of our mouse experimental unit (IERP, INRA, Jouy en Josas, France). We thank the @Bridge platform for use of the Agilent Bioanalyzer and for histology facilities (UMR 1313 GABI, Jouy-en-Josas, France), and particularly Marthe Vilotte for HE staining. We also thank the MIMA2 platform for providing access to the virtual slide scanner (Panoramic SCAN, 3DHISTECH). Finally, we are grateful to all members of the “Gonad Differentiation and its Disturbances” team for our scientific and technical discussions on Monday mornings, either in person or by videoconference. Vicky Hawken was responsible for the English language editing of this manuscript.

## Supporting information

**Supplementary Figure 1.** Validation of several DEGs by RT-qPCR (RNA-seq *Topaz1^−/−^ vs* WT testes). Validation of several differentially expressed up- or down-regulated genes and of non-DEGs from RNA-seq analysis by qRT-PCR from P16 (A) or P18 (B) mouse testis RNAs. The lines represent the median of each genotype (blue: WT; red: *Topaz1^−/−^*). A Kruskal-Wallis statistical test was performed (*p<0.05).

**Supplementary Figure 2.** Abnormal centrosome labeling in *Topaz1*-deficient gonads.

Immunofluorescence staining for γ-TUBULIN (red) and DAPI (blue) in WT (left) and *Topaz1^−/−^* (right) 30 d*pp* testes sections. Unlike the two red dots locating centrosomes in the meiotic metaphases seen in normal testes (left), centrosomes are abnormal in *Topaz1^−/−^* mutants (right) with one diffuse labeling. Zooms in white squares show spermatocytes in metaphase I (WT) or in metaphase I-like (*Topaz1^−/−^*) spermatocytes. Scale bar = 50μm.

**Supplementary Figure 3.** Reprogenomic data on the dynamic expression of *4930463O16Rik*. The dynamic expression of *4930463O16Rik* in five different tissues from male and female adult mice (A), and in embryonic primordial germ cells and adult male germ cells (B). *4930463O16Rik* is expressed in testes in germ cells during post-natal life. The strongest dynamic expression is found in pachytene spermatocytes.

**Supplementary Figure 4.** Reprogenomic data on the dynamic expression of *Gm21269*. The dynamic expression of *Gm21269* in five different tissues from male and female adult mice (A), in embryonic primordial germ cells and adult male germ cells (B). *Gm21269* is expressed in testes in germ cells during post-natal life. The strongest dynamic expression is found in pachytene spermatocytes.

**Supplementary Figure 5.** IGV representation of P18-testis RNA-seq. Expression of *4930463O16Rik* (A), *Gm21269* (B) and *4921513H07Rik* (C) from BigWig files of strand-specific RNA-seq data. The first four tracks represent transcripts of WT testes at P18; the next three tracks represent transcripts of *Topaz1^−/−^* testes at the same developmental stage. Representations of the genes (from mm10 or GRCm38) are shown at the bottom of each graph. A representation of the size of *4930463O16Rik* (A), *Gm21269* (B) and *492151H07Rik* (C) transcripts (red) from Ensembl data (GRCm38) is shown at the top. *4930463O16Rik* and *4921513H07Rik* gene transcriptions overlap in 3’ or 5’, respectively.

**Supplementary Figure 6.** Expression of *Gm21269*, *4930463O16Rik* and *492151H07Rik* mRNAs in testes from 5 days to adulthood. Quantitative RT-PCR analysis of *Gm21269*, *4930463O16Rik* and *492151H07Rik* gene expressions at different developmental stages in WT (blue) and *Topaz1^−/−^* (red) testes. The lines represent the median of each genotype. A Kruskal-Wallis statistical test was performed (*p<0.05; **p<0.01).

**Supplementary Figure 7.** *IS*H with PAS counterstained in WT mouse testes. Visualization of *4930463O16Rik* (A), *Gm21269* (B) and *4921513H07Rik* (C) mRNAs, respectively, by *IS*H at different seminiferous epithelium stages highlighted by PAS staining. Scale bar = 20μm

**Supplementary Figure 8.** LncRNA cellular localizations in testes from two month-old WT mice. *IS*H using (A) *4930463O16Rik*, (D) *Gm21269* and (G) *4921513H07Rik* probes (red). (B-E-H) Immunofluorescence staining with γH2Ax antibody was performed at the same stage of seminiferous epithelium to identify male germ cells (green). (C-F-I) DAPI (blue), visualizing nuclear chromosomes, was merged with *IS*H (green) and IF (red) signals. Zooms in white squares show spermatocytes during prophase I. No colocation between the sex body (γH2Ax) and lncRNAs (red) was evident. Scale bar = 20 μm.

**Supplementary Figure 9.** Validation of several DEGs by RT-qPCR (RNA-seq *4930463O16Rik^−/−^ vs* WT testes). Validation of several differentially expressed up- or down-regulated genes and of non-DEGs of RNA-seq analysis by RT-qPCR from P16 (A) or P18 (B) mouse testis RNAs. The lines represent the median of each genotype (blue: WT; red: 4930463O16Rik^−/−^). A Kruskal-Wallis statistical test was performed (*p<0.05).

**Supplementary Table 1.** List of DEGs in *Topaz1^−/−^* testes compared to WT. List of deregulated genes in *Topaz1* KO testes at P16 (sheet 1) and P18 (sheet 2) (adjusted p-value <0.05 and absolute Log2FC>1).

**Supplementary Table 2.** Functional annotation of P16 DEGs (RNA-seq *Topaz1^−/−^ vs* WT testes). DAVID functional Annotation Clustering (DAVID 6.8) analysis (based on GO terms and KEGG pathway) of all P16-differentially expressed genes (sheet 1) or only up-regulated DEGs (sheet 2) or down-regulated DEGs (sheet 3) in *Topaz1^−/−^* testes.

**Supplementary Table 3.** Functional annotation of P18 DEGs (RNA-seq *Topaz1^−/−^ vs* WT testes). DAVID functional Annotation Clustering (DAVID 6.8) analysis (based on GO terms and KEGG pathway) of P18-differentially expressed genes (sheet 1) or only up-regulated DEGs (sheet 2) or down-regulated DEGs (sheet 3) in *Topaz1^−/−^* testes. Annotation clusters based on the InterPro database of P18-down-regulated DEGs are mentioned in sheet 4.

**Supplementary Table 4.** List of primers. List of different primers used during this study for genotyping, RT-qPCR and gRNAs.

**Supplementary Table 5.** List of DEGs in *4930463O16Rik^−/−^* testes compared to WT. List of deregulated genes in *4930463O16Rik* KO testes at P16 (sheet 1) and P18 (sheet 2) (adjusted p-value <0.05 and absolute Log2FC>1).

**Supplementary Table 6.** Functional annotation of P18 DEGs (RNA-seq *4930463O16Rik^−/−^ vs* WT testes). DAVID functional Annotation Clustering (DAVID 6.8) analysis of P18-differentially expressed genes (sheet 1) or only up-regulated (sheet 2) or down-regulated DEGs (sheet 3) in *4930463O16Rik^−/−^* testes.

**Supplementary Table 7.** Casa system settings.

